# Shared genetic requirements for ATF5 translation in the vomeronasal organ and main olfactory epithelium

**DOI:** 10.1101/224980

**Authors:** Ryan P Dalton

**Affiliations:** Neuroscience Graduate Program, University of California, San Francisco, San Francisco, CA 94158, USA; Present address: The Miller Institute for Basic Research in Science, University of California, Berkeley, Berkeley, CA 94720, USA

## Abstract

**Background:** Both olfactory sensory neurons (OSNs) and vomeronasal sensory neurons (VSNs) require the transcription factor *Atf5* for maturation and survival. In OSNs, ATF5 translation is controlled by olfactory receptor (OR) expression-mediated activation of the PERK branch of the unfolded protein response. This study evaluated whether OSNs and VSNs share genetic requirements for ATF5 translation.

**Methods:** ATF5 immunoreactivity was assayed in whole vomeronasal organs from a series of genetic mutant animals identified in studies of OR gene choice, OR feedback, and regulation and OSN development.

**Results:** ATF5 expression in VSNs required the histone demethylase *Lsd1*, which has been previously reported to be required for OR expression. ATF5 expression also required PERK-mediated phosphorylation of the translation initiation factor eIF2α. Finally, unlike previous observations in OSNs, ATF5 was found to be widespread in the mature VNO and co-expressed with mature VSN markers.

**Conclusions:** These data suggest that the initiation of ATF5 translation in VSNs and OSNs is under similar regulation, and that persistent/prolonged ATF5 translation in VSNs may serve VSN-specific gene regulatory programs. This study firmly establishes the unfolded protein response as a major controller of sensory neuronal maturation and diversification.

## Introduction

Mice possess two olfactory organs: the main olfactory epithelium (MOE), which houses olfactory sensory neurons (OSNs), and the vomeronasal organ (VNO), which houses vomeronasal sensory neurons (VSNs)^1^. The MOE is thought largely to function in the detection of odors with no prior or innate behavioral importance, though there are notable exceptions^2-4 5^. The VNO, on the other hand, detects pheromones, which drive important social and reproductive behaviors^6,7^.

The VNO is a bilobal, cresecent-shaped neuroepithelium. It is neurogenic, giving rise to new VSNs throughout the life of the animal^1,8^. Immature VSNs are located at the tissue margins, or the tips of the crescents, while mature VSNs occupy more central areas. VSNs can be initially divided into two types, based on the expression of their primary signaling G proteins. Apical VSNs express the G protein *Gnai2*, as well as type I vomeronasal receptors (V1Rs). Basal VSNs express the G protein *Gnao* and type II vomeronasal receptors (V2Rs) ^9 10^. V1Rs are expressed monogenically and monoallelically^6,11,12^. The situation is markedly more complicated for V2Rs. Type II VSNs express two V2Rs in non-random combinations: one from V2R family A, B, or D; and at least one from V2R family C^13,14^. In addition, some basal VSNs also express at least one gene from the non-classical MHC *H2-Mv* gene family^15,16^.

The VR(s) expressed by a VSN both drive its pattern of connectivity to the accessory olfactory bulb and define its receptive field^12^. Therefore, the choice of receptor(s) to express is considered to be a central gene regulatory decision in VSN development. It is thought that the V1R expressed by a given type I VSN is chosen stochastically during development^1^. In the case of type II VSNs expressing multiple V2R genes, given that the coexpressed receptors (and *H2-Mv* genes for basal type II VSNs) occur in nonrandom combinations, it is possible that the initial V2R choice event is stochastic but acts to restrict subsequent V2R or *H2-Mv* choice events.

The past decade has seen the discovery of many molecular players Appendix (1) involved in the establishment of monogenic OR expression. Monogenic OR expression begins with OR gene choice, a complex process involving condensation of OSN chromatin^17^, extensive modification of the OR gene chromatin environment^18,19^, recruitment of *cis* and *trans* enhancer elements^20,21^, and cooperation between a number of transcriptional activators^22,23^. OR choice is followed by OR feedback, which functions to preclude further OR gene choice, to promote maturation of the OSN, and to stabilize expression of the chosen OR^1,24-29 30^ Together, OR choice and OR feedback ensure that each mature OSN expresses exactly one OR allele, defining each OSN as a sensitive and unambiguous signaling unit.

It was recently shown that ORs drive feedback by activating the unfolded protein response (UPR), a ubiquitous signaling pathway that homeostatically maintains the ER folding environment by modifying both its folding load and its folding capacity ^24,31^. OR expression activates the ER-resident kinase PERK, which then drives phosphorylation of the translation initiation factor eif2α, resulting in global attenuation of mRNA translation initiation and a specific increase in translation of mRNA encoding the transcription factor *Atf5*. ATF5 is required for OSN maturation and expression of adenylyl cyclase 3 (AC3). AC3 expression suppresses activity of a histone demethylase required for OR choice, LSD1. If OR choice fails to drive Adcy3 expression, as is the case with some OR pseudogenes as well as *Atf5* and *Adcy3* mutants, expression of the chosen OR is extinguished. This phenomenon, termed ‘gene switching’^27^, appears to add a layer of quality control for ORs, and also indicates that OR choice is initially unstable. This lack of stability is probably due to the dual demethylase activities of LSD1, which presumably allow it to de-silence an OR allele, and then to re-silence the same allele^19^. In this model, LSD1 downregulation by AC3 is required for stable transcription of the chosen OR. Together, these data support a model in which OR feedback acts to promotes OSN maturation, to prevent further OR choice, and to stabilize expression of the chosen OR allele.

In contrast to the growing body of knowledge on OR gene regulation, comparatively little is known for VRs. However, a number of lines of evidence support a model in which both V1Rs and V2Rs employ a feedback signal similar to that used by ORs to prevent further VR gene activation. First, VSNs choosing a V1R pseudogene target axons widely across the accessory olfactory bulb, indicating that they have subsequently selected a second V1R gene. This finding suggests that V1R protein activates vR feedback. Second, VSNs that choose an OR gene knocked into a V1R gene locus do not express additional V1Rs. This result suggests both that canonical V1R signaling is unimportant—as ORs and V1Rs signal through different second messengers—and that ORs can activate VR feedback^12^. Third, heterologous V2R expression activates the UPR, and both V1Rs and V2Rs, like ORs, fail to traffick from the ER when expressed heterologously. V2R trafficking appears to involve replacement of the ubiquitous chaperone Calreticulin with a VNO-specific homolog, Calreticulin 4. For V1Rs, the mechanism of ER trafficking in VSNs has yet to be established, but does not appear to involve either Calreticulin 4 or the OR transporters Rtp1/2^32^. Finally, it was recently shown that *Atf5* is required for maturation and survival of basal VSNs. This study also showed that while *Atf5* mRNA expression is ubiquitous in VSNs, ATF5 protein is expressed in more limited patterns, suggesting that *Atf5* mRNA is under translational control in the VNO^33^. In sum, these data suggest that OR and VR feedback may employ a common framework, converging on PERK-driven translation of ATF5.

In order to begin to define the mechanistic outline of VR feedback, I have assayed ATF5 protein expression in a series of mouse mutants previously employed in studies of OR feedback. I have found that in *Lsd1* mutant VNOs, ATF5 protein is absent, establishing a common genetic requirement for *Lsd1* in ATF5 translation in both the VNO and the MOE. Appearance of ATF5 also required both the ER-resident kinase PERK and phosphorylation of the translation initiation factor eif2α, suggesting that ER stress drives ATF5 translation in basal VSNs. Finally, in adult animals, ATF5 is widespread and found in anatomical areas corresponding to both immature and mature VSNs, suggesting that mature VSNs experience continued or spurious ER stress events. Together, these results support a model in which V1Rs and V2Rs both employ ER stress-mediated feedback, potentially with different requirements for ATF5 and subsequently with different transcriptional outcomes.

## Results

### *Lsd1* is required for ATF5 expression

*Lsd1* has previously been deleted from the olfactory placode by crossing animals carrying *loxP-surrounded Lsd1* alleles to animals expressing Cre recominase under the control of the FoxG1 promoter. These conditional mutants lose expression of most OR genes, resulting in a failure to translate ATF5 and a failure of OSNs to reach maturity^19^. To test whether VSNs and OSNs share a genetic requirement for *Lsd1* in the ATF5 expression, ATF5 immunoreactivity was assayed in control (*FoxG1-Cre; Lsd1 fl/+*) and mutant (*FoxG1-Cre; Lsd1 flfl*) VNO at embryonic day 18.5 (E18.5). A later analysis was not possible due to the perinatal lethality of this combination of alleles. As can be seen in Figure 1A-B, control animals exhibited robust ATF5 immunoreactivity in the VNO. As previously described, ATF5 expression was found to be widespread and heterogeneous from cell to cell. In contrast, *Lsd1* mutants did not have observable ATF5 expression (Figure 1C-D). Consistent with previous findings showing that *Atf5* is required for VSN maturation and survival, the VNO of the mutant animals was greatly reduced in size. Despite its decrease in size, the VNO was still readily identifiable through the use of a number of structural features, including the surrounding bone and mesenchyme structure, bilateral symmetry, position relative to the MOE, and the presence of a lumen of stereotypical shape, adjacent to an epithelium with a single layer of apical sustentacular cells. Together, these data indicate that in the VNO *Lsd1* is required for ATF5 expression, and by extension for VSN maturation. On the basis of these data, I hypothesize that VR expression is under *Lsd1* control, and that VR expression drives ATF5 translation. This hypothesis will be addressed in further detail in the discussion section.

**Figure 1:**
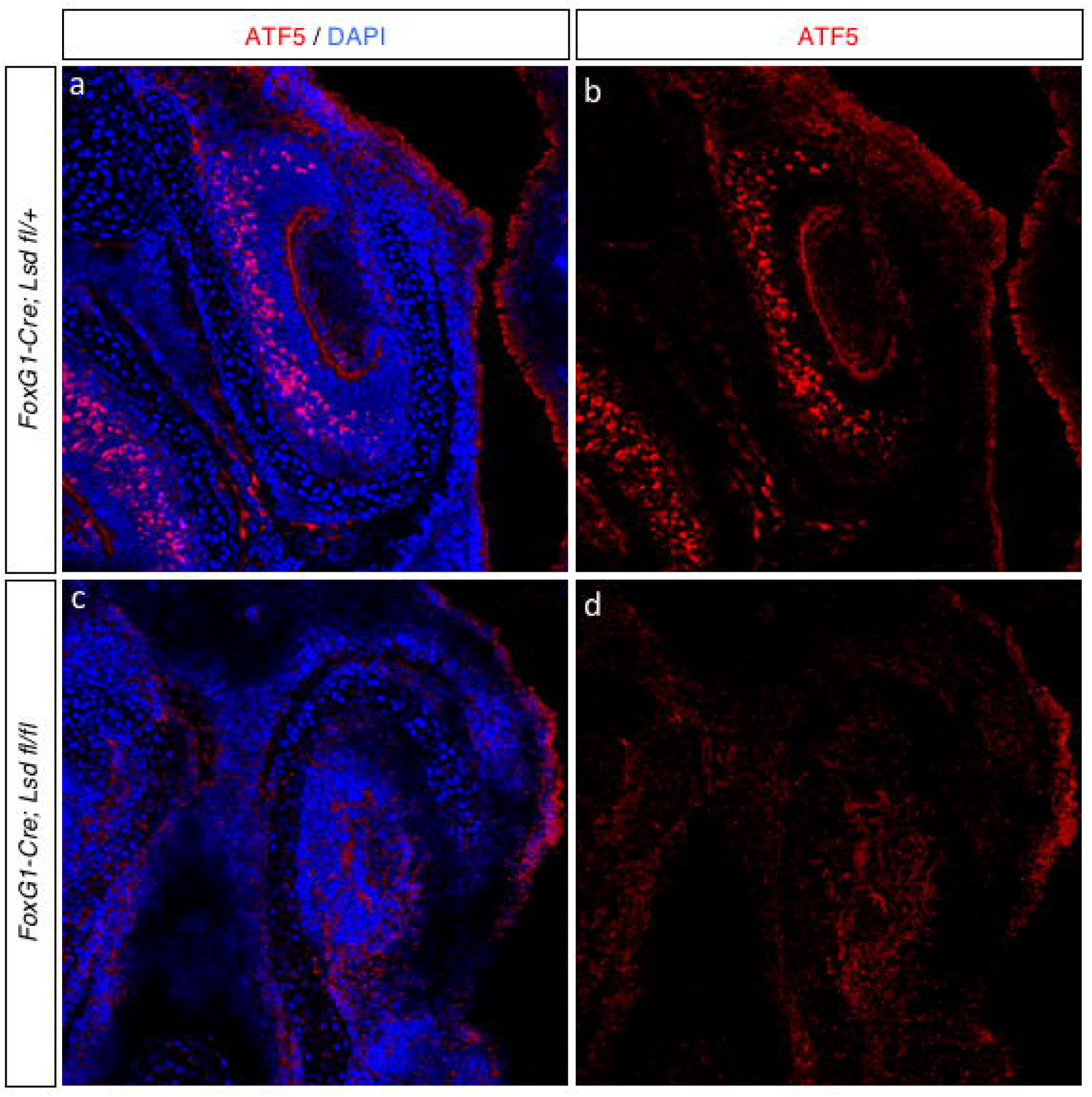
*Lsd1* is required for ATF5 expression. **(A-B)** Representative coronal section of embryonic day 18.5 (E18.5) VNO from a *FoxG1-Cre; Lsd1fl/+* animal. **(C-D)** Representative coronal section from a *FoxG1-Cre; Lsd1 fl/fl* VNO, also stained for ATF5 and DAPI. For all images, ATF5 immunoreactivity is shown in red and DAPI nuclear counterstain in blue.

### ATF5 expression requires PERK-mediated eif2α phosphorylation

I next asked whether ATF5 translation in the VNO is under the same regulatory control as in the MOE. The *Atf5* mRNA contains an inhibitory upstream open reading frame (iuORF) that under basal conditions suppresses its translation. However, upon phosphorylation of the translation initiation factor eIF2α at Serine-51, ribosomes bypass this iuORF to translate the *Atf5* mRNA coding sequence^24,34-37^. OR expression in the MOE promotes this phosphorylation event and ATF5 translation by activating the ER-resident kinase PERK. OR-driven ATF5 translation can be blocked either through PERK deletion or through mutation of the serine phosphorylation site on eIF2α to alanine. I therefore asked whether ATF5 was lost in the VNO of PERK mutants and eIF2α phosphor-mutants. While P0 *Perk*+/-VNO exhibited robust ATF5 immunoreactivity (Figure 2A-B), ATF5 was completely absent in littermate *Perk*-/-animals (Figure 2C-D). Similarly, ATF5 was completely absent in *eIF2tì^S51A/S51A^* animals, in which PERK is still present but cannot exert translational control through eif2α phosphorylation (Figure 2E-F). These data indicate that, as has been observed in the MOE and elsewhere, *Atf5* mRNA in the VNO is under translational regulation via PERK-dependent phosphorylation of eIF2α.

**Figure 2:**
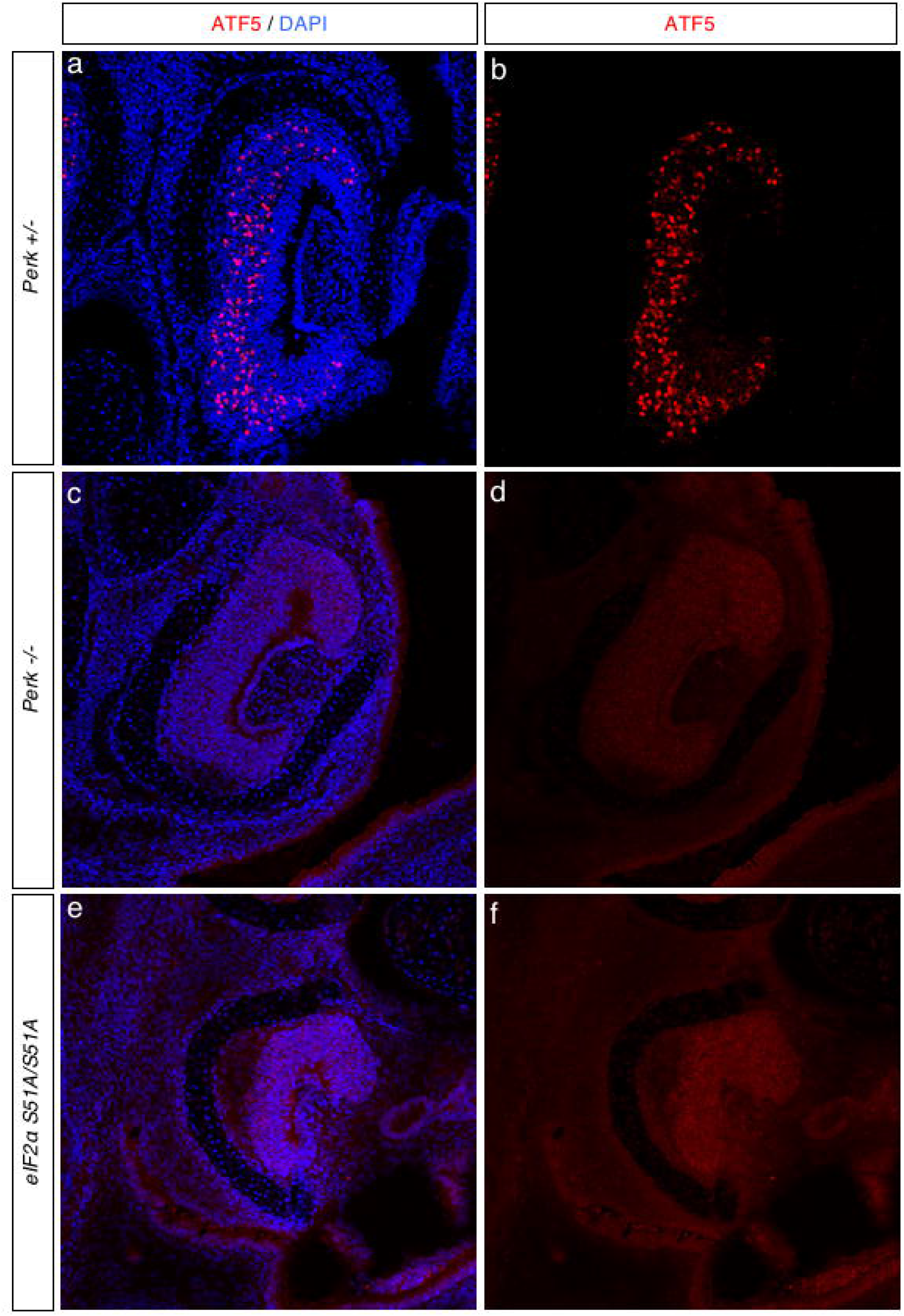
Translational control of ATF5. **(A-B)** Coronal section of postnatal day 0 (P0) VNO from *Perk+/-* animal. **(C-D)** Coronal section of P0 VNO from a *Perk-/-* littermate. **(E-F)** Coronal section from a PO *eIF2α*^S51A/S51A^ animal. For all images, ATF5 immunoreactivity is shown in red and DAPI nuclear counterstain in blue.

### ATF5 expression is widespread in adult animals

In the MOE, ATF5 expression is restricted to immature OSNs^24^. This expression pattern is intriguing, as both *Atf5* and OR mRNA continue to be expressed in mature OSNs. It has been proposed that this context-dependence for ATF5 translation is due to increased expression of OR transporters such as RTP1/2 in mature OSNs, which could compete ORs away from PERK or simply relieve the ER burden imposed by ORs. In the VNO, a previous report demonstrated that at P0, *Atf5* mRNA expression is essentially homogenous across the neuronal area, but that ATF5 protein is more heterogeneous. However, this report did not assay ATF5 expression in adult animals. While in young animals the VNO and MOE are dominated by immature neurons, in older animals the immature and mature neuronal compartments separate and resolve. In the adult VNO, a number of reports, using a variety of markers, have shown that immature VSNs are restricted to the VNO margins (i.e. the ‘tips’ of the VNO crescents) ^38,39^ Surprisingly, ATF5 was not restricted to the tissue margins in the adult VNO. Instead, it was widespread, heterogeneous, and found in areas corresponding to both immature and mature VSNs (Figure 3A-B). Furthermore, co-staining sections from this animal with antibody against olfactory marker protein (OMP) to label mature VSNs revealed that some ATF5-labeled cells co-express OMP. While most OMP+ cells were either ATF5-negative, or displayed barely-detectable ATF5, other OMP+ cells displayed saturating levels of ATF5. These observations suggest that, unlike in the MOE, in VSNs ATF5 continues to be expressed after maturation. Given the nature of ATF5 translational control and the consequences of persistent UPR activation, this raises a number of interesting questions regarding PERK activation dynamics and the transcriptional output of ATF5 in VSNs.

**Figure 3:**
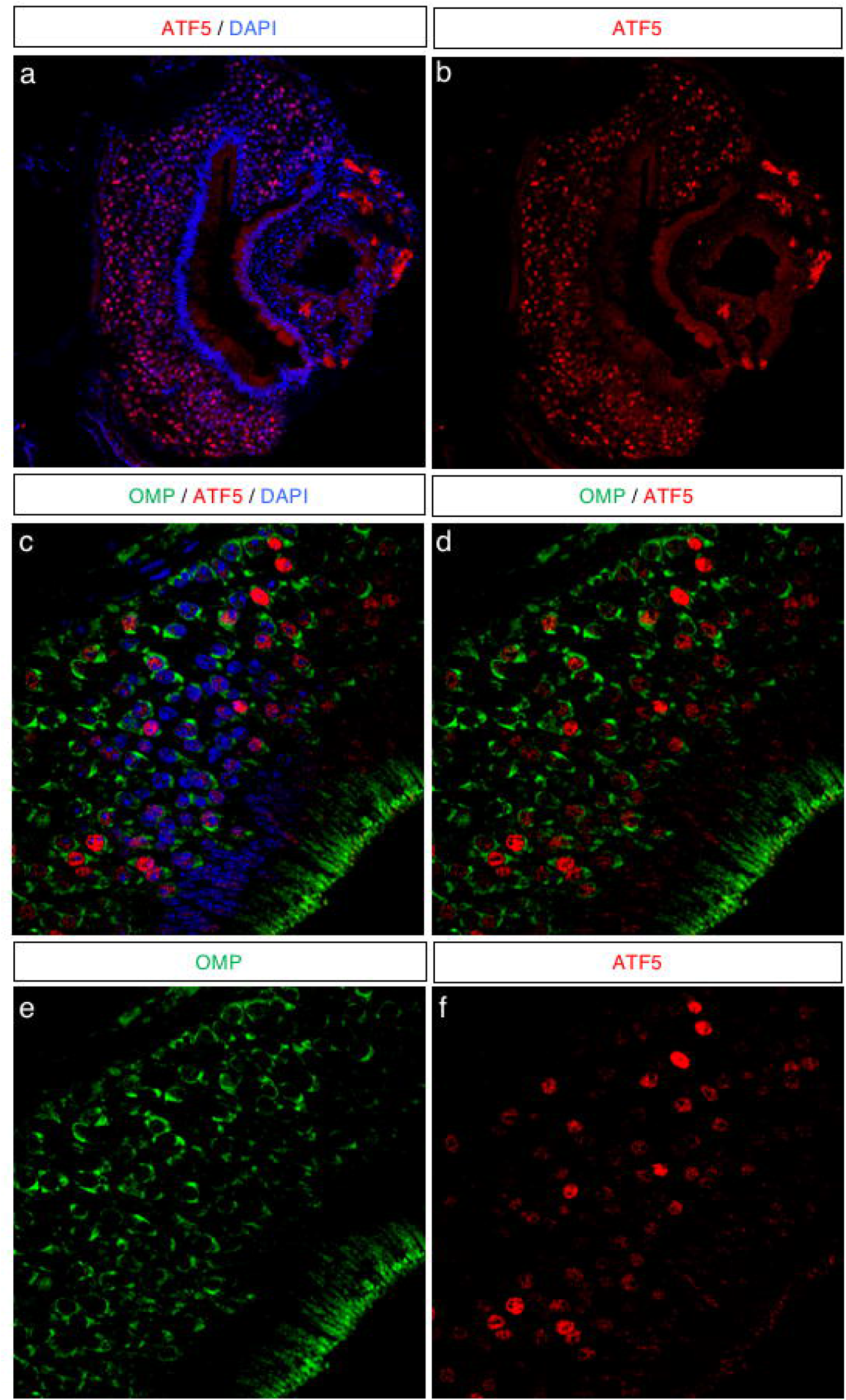
ATF5 expression in the adult VNO. **(A-B)** Coronal section of a postnatal day 35 VNO. ATF5 immunoreactivity is in red and DAPI in blue. **(C-F)** A close-up section from the same VNO as shown in A-B. Olfactory marker protein (OMP) immunoreactivity is shown in green, ATF5 in red, and DAPI in blue.

### Discussion

Receptor-driven feedback programs endow developing olfactory neurons with a means by which to establish distinct, unambiguous cell fates. These programs are therefore essential in the construction of the basic architecture of the olfactory system. Furthermore, because these feedback programs allow the appearance of a single protein to establish cell fate, they also act as an engine in neuronal diversification, forging a direct relationship between the number of chemoreceptor genes and the number of chemosensory cell fates.

My previous work uncovered that in OSNs, OR feedback is executed by co-option of the pERk branch of the unfolded protein response^24^. An obvious follow-up question to this work was whether this feedback mechanism was employed in other tissues in which sensory cells express single or small numbers of sensory receptors. In the present work, I demonstrate that expression of ATF5, which is required for OR feedback and for the maturation and survival of basal VSNs,^33^ has the same genetic requirements in the VNO and the MOE. This work therefore strongly suggests that ORs and VRs have a shared mechanism of feedback, converging on activation of the PERK branch of the UPR. This work was undertaken at the tissue level, and therefore more detailed analyses will likely be required in order to determine the specific requirements of VSN subtypes. Below I discuss some of the caveats of this work, as well as interesting questions for future work.

First, I hypothesized above that VR expression is under *Lsd1* control and is required for ATF5 expression. Several pieces of data prompted this hypothesis. Among them are the shared elements in VR and OR feedback discussed in the introduction such as their activation of the UPR in cell lines, as well as the requirement of *Lsd1* for ATF5 expression in the VNO demonstrated herein. However, it has yet to be directly demonstrated whether and how *Lsd1* influences VR expression, or whether VR expression in VSNs drives ATF5 translation. A number of experimental considerations make these analyses difficult. Among them are the prenatal lethality of *Lsd1* mutants and the requirement of *Atf5* for VSN survival^19,33^, which together result in exceedingly small amounts of tissue for analysis of VR expression or of the epigenetic landscape of the VR gene family. Additional genetic models would likely be useful in determining the role of *Lsd1* in VR choice, and a combination of biochemical and genetic approaches would be powerful in the determination of the mechanisms by which chemoreceptors influence PERK activity.

Second, it has been shown that a key element of OR feedback is AC3-driven downregulation of LSD1^19^. In the MOE, LSD1 downregulation both prevents further OR choice and acts to stabilize expression of the chosen OR. LSD1 downregulation therefore must be exquisitely timed. No analogous situation has yet been demonstrated for VSNs. It is worth noting that the requirements for VSNs are likely different than for OSNs. In particular, VSNs choosing VR pseudogenes continue to express them while also selecting another VR from a different VR gene cluster^40^. This finding indicates that VR choice may involve the permanent engagement of a single or limiting element in *cis* to a given VR cluster. VR feedback may therefore act to prevent further choice, but not to stabilize VR expression. Thus, if VSNs employ a mechanism similar to that of AC3 in OSNs, it may only act to terminate further VR choice, but not to stabilize VR choice. The mechanistic basis of this difference is a fascinating area for future study.

Third, the convergence on ATF5 in OSNs and VSNs prompts a number of questions on the role and transcriptional output of ATF5. For example, how could ATF5 control OR feedback in OSNs and VR feedback in VSNs? It seems likely, given that ATF5 is a bZip-family transcription factor, that ATF5 has different binding partners in different tissues. This model would allow for co-factors to tune the transcriptional specificity of ATF5, but would prevent their engagement until ATF5 has been translated. For example, in OSNs this may allow ORs to promote expression of RTP1/2 such that they can subsequently be targeted to the plasma membrane. In contrast, given that basal VSNs express non-random combinations of receptors and that the expression of these receptors is sequential, expression of one VR may drive ATF5 expression to aid in selection of a second VR (or an *H2-Mv*). The identity of these potential binding partners is a fascinating outstanding question and is likely to greatly aid in our understanding of chemoreceptor feedback programs.

Fourth, as demonstrated herein, ATF5 continues to be expressed in mature VSNs, unlike findings in OSNs. In addition, cell-to-cell levels of ATF5 appeared to be extremely variable, with signal nearly undetectable in most cells, but reaching saturation levels in other cells. This is a fascinating observation, as it would indicate that mature VSNs continue to experience ER stress events. *Atf5* is ubiquitous in VSNs^33^ and the UPR-driven mRNA translation program is rapidly induced but brief. I therefore hypothesize that the ATF5 expression patterns I observed reflect transient ER stress events experienced by many or all VSNs. However, it is impossible to rule out an alternate scenario in which some cells (or even VSN sub-types) experience continuing ER stress while others do not experience ER stress at all. An additional implication of the prolonged ATF5 expression pattern in VSNs could be that VRs and ORs have different mechanisms of PERK activation, for example direct versus indirect. A number of studies support an indirect model of PERK activation by ORs, in which ORs activate PERK only in the absence of RTP1/2, but this question is unaddressed for VRs. In addition, it is intriguing that ATF5 could be continuously expressed in mature VSNs, as it would beg the question of how VSNs can differentiate between bona fide ER stress and this developmental signal. Whether ATF5 has direct anti-apoptotic functions in VSNs as has been observed in other cell types has yet to be determined.

Finally, these findings firmly establish that a pathway canonically thought to be involved in the detection and resolution of cellular stress responses is fundamental in the designation of cellular identity and in cell maturation. This not only begs a reassessment of the role of PERK signaling, but also suggests specific additional studies. Given that activation of PERK provides such a powerful means by which to coordinate receptor appearance to the cellular gene expression program, and given that a multitude of cell types are defined by their expression of one or a handful of receptors, it would be surprising if PERK were not involved in other receptor-driven feedback programs. Excellent candidates include somatosensory neurons expressing Mas-related GPR family members, taste receptor cells, and photoreceptor cells. Specific chaperone or transporter requirements for these different receptors would provide a simple and generalizable mode for receptors to activate PERK in order to drive global gene expression programs, whose outputs can then be tuned by the use of tissue or cell type-specific co-factors.

## Methods

### Mice and strains used

All mice were housed in standard conditions with a 12-hour light/dark cycle and access to food and water in a UCSF barrier facility. All mouse experiments as well as euthanasia were approved by and were in accordance with University of California, San Francisco Institutional Animal Care and Use Committee (IACUC) protocol as described previously^18^. Animals used in this study were under protocols held by the Lomvardas laboratory. Details on standard procedures including euthanasia can be found at the UCSF IACUC website (http://iacuc.ucsf.edu/Policies/awStandardProcedures.asp). Because all animals described in this study were only used for tissue collection, the relevant UCSF IACUC setions are those that deal with proper euthanasia. For all animals used in this study, animals were single or pair-bred (for animals harvested during pregnancy) or were group housed (for animals harvested as adults). For prenatal experiments, pregnancies were timed such that pregnant females and pups could be harvested at the time points listed in the study. The age at which tissue was collected for each experiment is indicated in the figure legends. For all tissue collected, animals were euthanized at the indicated time point using a CO2 flow rate designed to minimize suffering and per IACUC regulations. Following CO2 exposure, animals were secondarily euthanized either by cervical dislocation for adults or decapitation for prenatal and perinatal pups, and then immediately dissected. Main olfactory epithelia or vomeronasal organs were dissected using clean, sterile, and recently sharpened tools. All strains were maintained on a mixed (Black 6 / FVB) genetic background. The following mouse lines used in this study have been previously described: *FoxG1-Cre; Lsd1 flox*^19^, *Perk* and *Eif2αS51A/S51A^24^*. No sex-based differences in expression of the markers tested herein were identified in preliminary studies and the studies herein contained animals of both sexes. All efforts were made to minimize the number of animals used in performing this study.

### Immunofluorescence

Immunofluorescence (IF) was performed as previously described^17,19,24^. All animals were dissected immediately following euthanasia by CO2 exposure. Briefly, tissue was directly dissected into optimal cutting temperature compound (OCT, Tissue-Tek #4583). 14μm sections were air-dried on glass slides (VWR #48311-6703) for 10 minutes, fixed in 4% paraformaldehyde (PFA, Sigma #158127) in phosphate buffer solution (PBS) for 10 minutes, washed 3x5 minutes in PBS + .1% Triton-X (PBST), blocked for 1 hour in 4% donkey serum in PBST, then incubated with primary antibodies diluted in PBST and under coverslips (VWR # 470019-008) overnight at 4C. The following day, slides were washed 3x15 minutes in PBST and then incubated with secondary antibodies and DAPI in PBST at concentrations of 1:1000 under cover slides. Slides were then washed 3x15 minutes in PBST and mounted with vectashield (Vector Laboratories # H-1000) for imaging. Imaging was performed on Leica 700-series laser scanning confocal microscopes. The following antibodies were used: goat anti-Atf5 (Santa Cruz Biotechnology, SC-46934, dilution 1:250), rabbit anti-OMP (Abcam ab93127, dilution 1:250). For each panel, at least one mouse per genotype was sectioned. Differences were not noted between males and females. Mice were genotyped in-house using genotyping protocols suggested by the original generators of the mouse line. Genotyping protocols included positive and negative controls reactions. Full MOE or VNO were sectioned. To minimize variability between slides, control and experimental genotypes were sectioned onto the same slide. Slides with strong antibody signal and low background were selected for analysis. For each section, microscope settings were optimized for signal:noise. All image analysis was done in Fiji and consisted only of changing brightness and contrast for each channel.

### Amelioration of animal suffering

All efforts were made to ameliorate the suffering of animals used in this study. Animals received regular care from the author and from the animal facility personnel, as well as monitoring for health and injury. Unhealthy or injured animals were either treated or, in severe cases, humanely euthanized. In accordance with UCSF IACUC guidelines (see above), all animals were euthanized by exposure to CO2 at a rate deemed to minimize stress and suffering. Animals were not bred unnecessarily, and when possible multiple types of tissue were dissected from each animal and saved for potential future use.

### Data availability

Materials and data are available upon reasonable request from the author. Raw image files are available through F1000 Research.

### Grant information

This work was funded by a Genentech Predoctoral Fellowship. RPD conceived and executed experiments and wrote this manuscript.

## References

1. Dalton, R. P. & Lomvardas, S. Chemosensory receptor specificity and regulation. Annu. Rev. Neurosci. 38, 331–349 (2015).

2. Buck, L. & Axel, R. A novel multigene family may encode odorant receptors: a molecular basis for odor recognition. Cell 65, 175–187 (1991).

3. Liberles, S. D. & Buck, L. B. A second class of chemosensory receptors in the olfactory epithelium. Nature 442, 645–650 (2006).

4. Omura, M. & Mombaerts, P. Trpc2-expressing sensory neurons in the mouse main olfactory epithelium of type B express the soluble guanylate cyclase Gucy1b2. Mol. Cell. Neurosci. 65, 114–124 (2015).

5. Greer, P. L. et al. A Family of non-GPCR Chemosensors Defines an Alternative Logic for Mammalian Olfaction. Cell 165, 1734–1748 (2016).

6. Dulac, C. & Axel, R. A novel family of genes encoding putative pheromone receptors in mammals. Cell 83, 195–206 (1995).

7. Halpern, M. The Organization And Function Of The Vomeronasal System. Annu. Rev. Neurosci. 10, 325–362 (1987).

8. Brennan, P. & Keverne, E. in Handbook of Olfaction and Gustation (CRC Press, 2009). doi: 10.1201/9780203911457.ch46

9. Berghard, A. & Buck, L. B. Sensory transduction in vomeronasal neurons: evidence for G alpha o, G alpha i2, and adenylyl cyclase II as major components of a pheromone signaling cascade. Journal of Neuroscience 16, 909–918 (1996).

10. Jia, C. & Halpern, M. Subclasses of vomeronasal receptor neurons: differential expression of G proteins (Gi 2 and Go) and segregated projections to the accessory olfactory bulb. Brain Research 719, 117–128 (1996).

11. Ryba, N. J. & Tirindelli, R. A new multigene family of putative pheromone receptors. Neuron 19, 371–379 (1997).

12. Rodriguez, I., Feinstein, P. & Mombaerts, P. Variable patterns of axonal projections of sensory neurons in the mouse vomeronasal system. Cell 97, 199–208 (1999).

13. Martini, S., Silvotti, L., Shirazi, A., Ryba, N. J. & Tirindelli, R. Co-expression of putative pheromone receptors in the sensory neurons of the vomeronasal organ. J. Neurosci. 21, 843–848 (2001).

14. Ishii, T. & Mombaerts, P. Coordinated coexpression of two vomeronasal receptor V2R genes per neuron in the mouse. Molecular and Cellular Neuroscience 46, 397–408 (2011).

15. Ishii, T. & Mombaerts, P. Expression of Nonclassical Class I Major Histocompatibility Genes Defines a Tripartite Organization of the Mouse Vomeronasal System. Journal of Neuroscience 28, 2332–2341 (2008).

16. Ishii, T., Hirota, J. & Mombaerts, P. Combinatorial Coexpression of Neural and Immune Multigene Families in Mouse Vomeronasal Sensory Neurons. Current Biology 13, 394–400 (2003).

17. Clowney, E. J. et al. Nuclear aggregation of olfactory receptor genes governs their monogenic expression. Cell 151, 724–737 (2012).

18. Magklara, A. et al. An epigenetic signature for monoallelic olfactory receptor expression. Cell 145, 555–570 (2011).

19. Lyons, D. B. et al. An epigenetic trap stabilizes singular olfactory receptor expression. Cell 154, 325–336 (2013).

20. Lomvardas, S. et al. Interchromosomal Interactions and Olfactory Receptor Choice. Cell 126, 403–413 (2006).

21. Markenscoff-Papadimitriou, E. et al. Enhancer interaction networks as a means for singular olfactory receptor expression. Cell 159, 543–557 (2014).

22. Monahan, K. et al. Cooperative interactions enable singular olfactory receptor expression in mouse olfactory neurons. Elife 6, 1083 (2017).

23. Monahan, K. & Lomvardas, S. Monoallelic expression of olfactory receptors. Annu. Rev. Cell Dev. Biol. 31, 721–740 (2015).

24. Dalton, R. P., Lyons, D. B. & Lomvardas, S. Co-opting the unfolded protein response to elicit olfactory receptor feedback. Cell 155, 321–332 (2013).

25. Sharma, R. et al. Olfactory receptor accessory proteins play crucial roles in receptor function and gene choice. Elife 6, 1083 (2017).

26. Lewcock, J. W. & Reed, R. R. A feedback mechanism regulates monoallelic odorant receptor expression. … Academy of Sciences of the United … (2004).

27. Shykind, B. M. et al. Gene switching and the stability of odorant receptor gene choice. Cell 117, 801–815 (2004).

28. Serizawa, S. et al. Negative Feedback Regulation Ensures the One Receptor-One Olfactory Neuron Rule in Mouse. Science 302, 2088–2094 (2003).

29. Nguyen, M. Q., Zhou, Z., Marks, C. A., Ryba, N. J. P. & Belluscio, L. Prominent Roles for Odorant Receptor Coding Sequences in Allelic Exclusion. Cell 131, 1009–1017 (2007).

30. Abdus-Saboor, I. et al. An Expression Refinement Process Ensures Singular Odorant Receptor Gene Choice. Curr. Biol. 26, 1083–1090 (2016).

31. Walter, P. & Ron, D. The unfolded protein response: from stress pathway to homeostatic regulation. Science 334, 1081–1086 (2011).

32. Dey, S. & Matsunami, H. Calreticulin chaperones regulate functional expression of vomeronasal type 2 pheromone receptors. Proc. Natl. Acad. Sci. U.S.A. 108, 16651–16656 (2011).

33. Nakano, H. et al. Activating transcription factor 5 (ATF5) is essential for the maturation and survival of mouse basal vomeronasal sensory neurons. Cell Tissue Res. 363, 621–633 (2016).

34. Hansen, M. B. et al. Mouse Atf5: molecular cloning of two novel mRNAs, genomic organization, and odorant sensory neuron localization. Genomics 80, 344–350 (2002).

35. Zhou, D. et al. Phosphorylation of eIF2 directs ATF5 translational control in response to diverse stress conditions. J. Biol. Chem. 283, 7064–7073 (2008).

36. Watatani, Y. et al. Stress-induced translation of ATF5 mRNA is regulated by the 5’-untranslated region. J. Biol. Chem. 283, 2543–2553 (2008).

37. Greene, L. A., Lee, H. Y. & Angelastro, J. M. The transcription factor ATF5: role in neurodevelopment and neural tumors. J. Neurochem. 108, 11–22 (2009).

38. Oboti, L. et al. Pregnancy and estrogen enhance neural progenitor-cell proliferation in the vomeronasal sensory epithelium. BMC Biol. 13, 104 (2015).

39. Enomoto, T. et al. Bcl11b/Ctip2 controls the differentiation of vomeronasal sensory neurons in mice. J. Neurosci. 31, 10159–10173 (2011).

40. Roppolo, D. et al. Gene cluster lock after pheromone receptor gene choice. EMBO J. 26, 3423–3430 (2007).

